# Innate immune response to SARS-CoV-2 infection contributes to neuronal damage in human iPSC-derived peripheral neurons

**DOI:** 10.1101/2022.11.18.517047

**Authors:** Vania Passos, Lisa M. Henkel, Jiayi Wang, Francisco J. Zapatero-Belinchón, Rebecca Möller, Guorong Sun, Inken Waltl, Birgit Ritter, Kai A. Kropp, Shuyong Zhu, Michela Deleidi, Ulrich Kalinke, Günter Höglinger, Gisa Gerold, Florian Wegner, Abel Viejo-Borbolla

## Abstract

Severe acute respiratory coronavirus 2 (SARS-CoV-2) infection causes neurological disease in some patients suggesting that infection can affect both the peripheral and central nervous system (PNS and CNS, respectively). It is not clear whether the outcome of SARS-CoV-2 infection of PNS and CNS neurons is similar, and which are the key factors that cause neurological disease: SARS-CoV-2 infection or the subsequent immune response. Here, we addressed these questions by infecting human induced-pluripotent stem cell-derived CNS and PNS neurons with the β strain of SARS-CoV-2. Our results show that SARS-CoV-2 infects PNS neurons more efficiently than CNS neurons, despite lower expression levels of angiotensin converting enzyme 2. Infected PNS neurons produced interferon λ1, several interferon stimulated genes and proinflammatory cytokines. They also displayed neurodegenerative-like alterations, as indicated by increased levels of sterile alpha and Toll/interleukin receptor motif-containing protein 1, amyloid precursor protein and α-synuclein and lower levels of nicotinamide mononucleotide adenylyltransferase 2 and β-III-tubulin. Interestingly, blockade of the Janus kinase and signal transducer and activator of transcription pathway by Ruxolitinib did not increase SARS-CoV-2 infection, but reduced neurodegeneration, suggesting that an exacerbated neuronal innate immune response contributes to pathogenesis in the PNS.

## Introduction

SARS-CoV-2 is the causative agent of coronavirus disease 2019 (COVID-19). Clinical studies indicate that some patients infected with SARS-CoV-2 develop neurological disease in the central and peripheral nervous systems (CNS and PNS, respectively) (Alipoor et al., 2020; Helms et al., 2020; Mao et al., 2020; Paterson et al., 2020; Politi et al., 2020; Taquet et al., 2021; Zanin et al., 2020). Diseases associated with the PNS occurring after SARS-CoV-2 infection include nerve pain, Guillain-Barré syndrome, myasthenia gravis, neurosensory disorders and peripheral neuropathy (Abdelnour et al., 2020; Andalib et al., 2021; Huber et al., 2020; Mao et al., 2020; McFarland et al., 2021; Romero-Sanchez et al., 2020; Zhao et al., 2020). Moreover, many individuals with long COVID-19 suffer from neurological disorders including brain fog and cognitive impairment, implying the persistence of neurological damage or even progression to neurodegenerative disease (Ali et al., 2022; Alipoor et al., 2020; Spudich and Nath, 2022; Stefanou et al., 2022). Interestingly, some long COVID-19 patients with neurological symptoms have increased loss of small nerve fibres in the cornea (Bitirgen et al., 2021) and peripheral neuropathy, probably due to an exacerbated immune response to infection (Oaklander et al., 2022).

Currently, it is not clear whether neurological diseases in COVID-19 patients are due to direct effects of viral infection of the nervous system or to an indirect impact of the immune response. SARS-CoV-2 RNA genome, transcripts and antigens were detected in the cerebrospinal fluid (CSF) and brains of some, but not all, COVID-19 patients that developed neurological abnormalities following infection (Farhadian et al., 2020; Moriguchi et al., 2020; Puelles et al., 2020; Solomon et al., 2020; Song et al., 2021; Wu et al., 2020). It has been suggested that a dysregulation of the immune response, including the induction of cytokine storm, is responsible for neurological complications during COVID-19 (Andalib et al., 2021; Koenigsknecht-Talboo and Landreth, 2005; Magliozzi et al., 2018; Nagu et al., 2021; Pilotto et al., 2021; Schwarzschild et al., 2008).

To investigate the underlying mechanisms of neurological symptoms following SARS-CoV-2 infection, several groups employed human neurons and neuronal models derived from stem cells to study SARS-CoV-2 infection. Most of the studies focused on neurons of the CNS rather than on those of the PNS. SARS-CoV-2 infects human iPSC-derived CNS neurons and brain organoids (Bauer et al., 2021; Bullen et al., 2020; Jacob et al., 2020; McMahon et al., 2021; Pellegrini et al., 2020; Ramani et al., 2020; Song et al., 2021; Tiwari et al., 2021). In one of the few studies addressing SARS-CoV-2 infection of human PNS neurons, the virus efficiently infected human embryonic stem cells-derived peripheral neurons and affected the expression of chemosensory genes (Lyoo et al., 2022).

Neurons can detect viruses through pattern recognition receptors and express and respond to interferon (IFN) to combat virus infection (Cavanaugh et al., 2015; Chhatbar et al., 2018; Delhaye et al., 2006; Gao et al., 2021; Ghita et al., 2021; Li et al., 2011; Tiwari et al., 2021; Zhou et al., 2009). Neurons also respond to viral infections through induction of axonal degeneration, apoptosis and autophagy, although the latter can also be proviral (Chawla et al., 2022; Crawford et al., 2022; Mori et al., 2004; Sundaramoorthy et al., 2020; Tsunoda, 2008; Yordy et al., 2012). Under certain circumstances, the innate and intrinsic neuronal responses to viral infection can lead to neurodegenerative processes, e.g., by induction of the unfolded protein response (UPR) (Alfano et al., 2019; Medigeshi et al., 2007) and activation of sterile alpha and Toll/interleukin receptor motif-containing protein 1 (SARM1), a protein that triggers neurite degeneration (Gerdts et al., 2015; Hou et al., 2013; Osterloh et al., 2012; Sundaramoorthy et al., 2020).

Here, we addressed the impact of SARS-CoV-2 infection and the subsequent innate and intrinsic immune responses on the induction of neuronal damage using human induced pluripotent stem cell (iPSC)-derived CNS and PNS neurons. Both cell types expressed very low mRNA levels of angiotensin-converting enzyme 2 (ACE2), a known SARS-CoV-2 receptor, and other entry factors. Interestingly, neurons from the PNS were much more permissive to SARS-CoV-2 than those from the CNS. This was accompanied by increased type III IFN response, loss of cytoskeleton proteins (β-III-tubulin and microtubule-associated protein 2, MAP2), as well as gene and protein expression profiles characteristic of neurite degeneration. Interestingly, inhibition of the Janus kinase and signal transducer and activator of transcription (JAK/STAT) pathway by Ruxolitinib reduced neurite degeneration, while having no significant effect on SARS-CoV-2 infection. Our results suggest that there are important differences in the way that CNS and PNS neurons respond to SARS-CoV-2 infection and link the innate immune response with neurodegeneration in human PNS neurons.

## Results

### iPSC-derived human CNS and PNS neurons express low levels of SARS-CoV-2 receptors and entry factors

To perform a comparative analysis of SARS-CoV-2 infection in PNS and CNS neurons, we employed protocols developed in our laboratories to differentiate human iPSC into neurons with characteristics of sensory and striatal medium spiny neurons (MSN) (schematic depiction in Figure 1A and 1B) (Kutschenko et al., 2021; Zhu et al., 2021). We will refer to the sensory neurons and MSN as PNS and CNS neurons, respectively. SARS-CoV-2 cell entry normally requires processing of the S protein by transmembrane protease, serine 2 (TMPRSS2), followed by binding to ACE2 (Hoffmann et al., 2020; Zhou et al., 2020). Neuropilin-1 (Nrp-1) seems to facilitate SARS-CoV-2 infection (Cantuti-Castelvetri et al., 2020; Daly et al., 2020) and mediates entry into human astrocytes in brain organoids (Kong et al., 2022). We determined the mRNA levels of ACE2, TMPRSS2 and Nrp-1 using quantitative PCR. Both neuronal subtypes expressed low mRNA levels of the analysed genes and no detectable ACE2 protein, while Vero76 cells, used to prepare SARS-CoV-2 stocks, expressed higher ACE2 mRNA and protein (Figure 1C,D).

**Figure 1:**
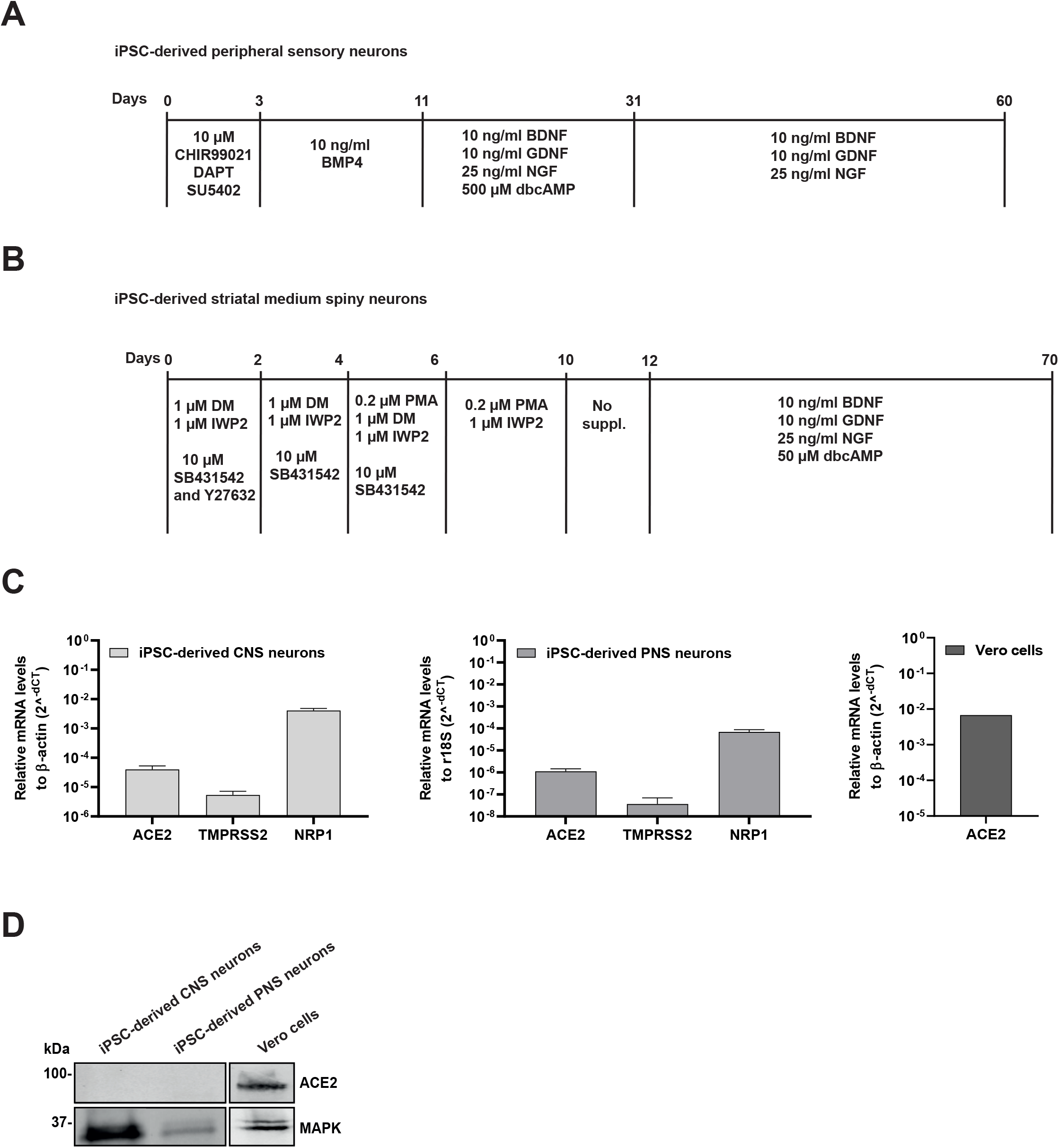
Differentiation of human iPSC-derived CNS and PNS neurons, and their expression profiles of SARS-CoV-2 receptors and entry factors. (A,B) PNS (A) and CNS (B) neurons were differentiated from iPSC adding the indicated supplements at the corresponding time points (Kutschenko et al., 2021; Zhu et al., 2021). (C) Graphs showing relative expression of *ACE2, TMPRSS2* and *NRP1* in iPSC-derived CNS neurons (left) and PNS neurons (middle) and that of *ACE2* in Vero76 cells (right). Gene expression was set relative to β-actin or 18S. Error bars represent standard deviation of the arithmetic mean from three independent experiments. (D) Western blot showing detection of ACE2 and MAPK in cell lysates of iPSC-derived CNS and PNS neurons and Vero76 cells. Shown is one representative blot out of three independent ones. Abbreviations: iPSC, induced pluripotent stem cells; DAPT, γ-secretase inhibitor; BMP4, bone morphogenetic protein 4; BDNF, brain-derived neurotrophic factor; GDNF, glial cell line-derived neurotrophic factor; NGF, nerve growth factor; dbcAMP: dibutyryl cyclic adenosine monophosphate; PMA, purmorphamine; DM, dorsomorphin; IWP2, Wnt antagonist 2; SB431542, TGF-β inhibitor; Y27632, Rock inhibitor.

### SARS-CoV-2 infects a low number of iPSC-derived human CNS neurons and does not trigger a strong innate immune response

We addressed whether SARS-CoV-2 could infect iPSC-derived human CNS neurons despite the low expression level of known SARS-CoV-2 entry factors. Infection of CNS neurons with SARS-CoV-2 β strain at a multiplicity of infection (MOI) of 0.01 led to low expression of nucleocapsid (NC) protein compared to the results obtained with infected Vero76 cells (Figure 2A and Supplementary Figure 1A). We could not detect NC protein in Vero76 cells infected with UV-inactivated SARS-CoV-2, suggesting that the detected protein corresponded to newly expressed NC (Supplementary Figure 1A). We also detected *NC* and *Membrane* transcripts in CNS neurons infected with SARS-CoV-2 (Supplementary Figure 1B). To address whether there was productive infection of CNS neurons, we performed a kinetic experiment and observed that SARS-CoV-2 mRNA expression increased in a time and MOI dependent manner (Supplementary Figure 1B). These results suggested that SARS-CoV-2 replicated in CNS neurons, although with low efficiency. To address whether the neuronal innate immune response could limit virus replication, we blocked the JAK/STAT pathway with Ruxolitinib prior to exposure of CNS neurons to SARS-CoV-2. Interestingly, Ruxolitinib did not modulate the infection efficiency of SARS-CoV-2 at the translational or transcriptional level (Figure 2A,B).

**Figure 2:**
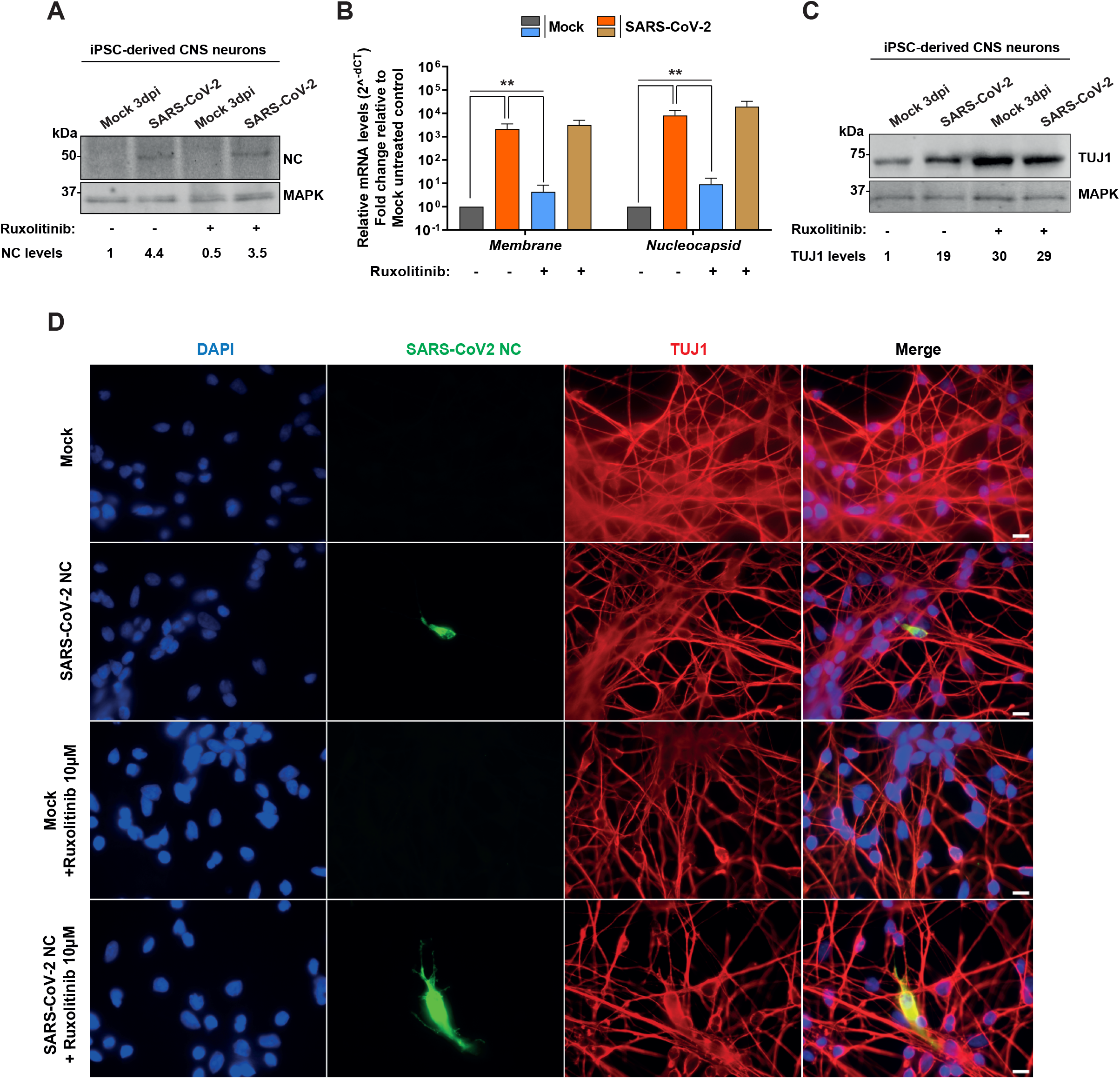
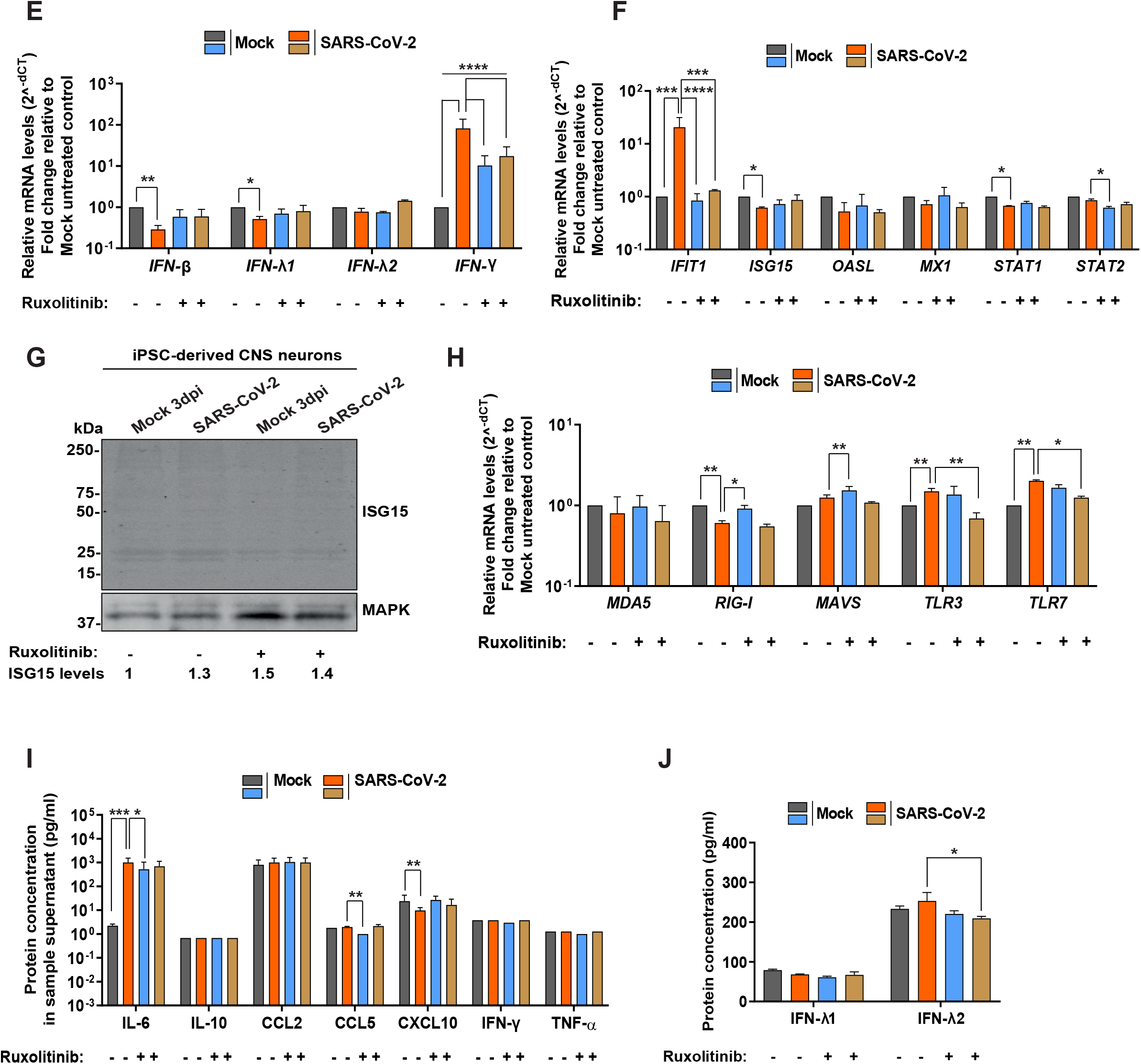
SARS-CoV-2 infects CNS neurons with low efficiency without eliciting a robust IFN and ISG response. (A) Western blot showing SARS-CoV-2 NC and MAPK in CNS neurons lysed at 3 dpi. NC levels relative to MAPK were measured with ImageJ. Shown is a representative blot from 3 independent experiments. (B) Graphs showing relative expression of SARS-CoV-2 *Membrane* and *Nucleocapsid* mRNA in CNS neurons at 3 dpi. (C) Immunoblot showing TUJ1 and MAPK in cell lysates of CNS neurons at 3 dpi. TUJ1 levels relative to MAPK were measured using ImageJ. Shown is one representative blot out of three independent ones. (D) Immunofluorescence of CNS neurons fixed at 3 dpi. The cells were stained with anti-SARS-CoV-2 NC, anti-TUJ1 and DAPI. Scale bar = 20 μm; Amplification: 100 X. (E, F, H) Graphs showing relative expression of host genes in CNS neurons at 3 dpi. (G) Immunoblot to detect ISG15 and MAPK in cell lysates of CNS neurons. Relative ISG15 levels were measured using ImageJ. Shown is a representative blot out of three independent ones. (I,J) Graph showing protein concentrations of several cytokines in supernatants obtained from CNS neurons at 3 pdi determined by cytometric bead array multiplex analysis (I) and ELISA (J). In all immunoblots, the numbers below the blot represent the fold change to mock-infected, untreated control. In all graphs showing gene expression, this was quantified by RT-qPCR and set relative to β-actin. Fold-change is relative to mock-infected, untreated control. In all graphs, error bars represent standard deviation of the arithmetic mean from three independent experiments. Statistical analyses for all experiments were determined using one-way ANOVA followed by the Dunnetts multiple comparison post-test. \**P*<0.03; \*\**P*<0.002; \*\*\**P*<0.0002; \*\*\*\**P*<0.0001; not significant comparisons are not indicated.

Co-staining of NC protein and β-III-tubulin (TUJ1) indicated that neurons were infected (Figure 2D). The number of NC positive CNS neurons was low and dispersed in the culture. We did not detect clear cytopathic effect and the protein level of the cytoskeleton protein β-III-tubulin did not decrease upon SARS-CoV-2 infection (Figure 2C). Instead, we observed more β-III-tubulin compared to the mock-infected cells, suggesting no degradation of this cytoskeletal protein upon infection of CNS neurons. Blocking the JAK/STAT pathway resulted in higher β-III-tubulin level in CNS neurons that were either mock- or SARS-CoV-2-infected (Figure 2C). Overall, these results show that SARS-CoV-2 can infect a limited number of iPSC-derived human CNS neurons, which do not appear to be severely damaged by the presence of the virus. We then addressed whether CNS neurons responded to infection with SARS-CoV-2 by expressing IFN and interferon stimulated genes (ISGs). Infection with SARS-CoV-2 reduced the expression of type I (*IFN-β*) and type III IFN (*IFN-λ1*) and increased that of *IFN-γ*, but IFN protein levels were not increased (Figure 2I,J and Supplementary Figure 2A). Infection decreased the expression of *ISG15* and *STAT1*, induced the expression of interferon induced protein with tetratricopeptide repeats 1 (*IFIT1*) and did not modify that of 2’-5’-Oligoadenylate synthetase like (*OASL*), interferon-induced GTP-binding protein (*Mx1*), and *STAT2* (Figure 2F,G). Ruxolitinib significantly decreased the expression of *IFN-γ* and *IFIT1*, indicating that the treatment was effective (Figure 2E,F). These results indicate that infection of CNS neurons with SARS-CoV-2 did not induce a strong expression of type I and III IFN and ISGs.

We then investigated the expression level of pattern recognition receptors known to be involved in SARS-CoV-2 recognition, namely melanoma differentiation-associated protein 5 (MDA-5), retinoic acid-inducible gene I (RIG-I), mitochondrial antiviral-signalling protein (MAVS) and Toll-like receptor 3 and 7 (TLR3 and TLR7, respectively). Interestingly, *RIG-I* was decreased in the presence of SARS-CoV-2, while the expression of *TLR3* and *TLR7* increased upon infection (Figure 2H). Addition of Ruxolitinib during infection reduced the expression level of *TLR3* and *TLR7*.

We then determined the level of cytokines present in the supernatant of mock- and SARS-CoV-2-infected CNS neurons in the absence and presence of Ruxolitinib. The levels of CXCL10 were lower in infected cells and, with the exception of interleukin 6 (IL-6), none of the analysed cytokines were increased during SARS-CoV-2 infection (Figure 2I), suggesting that CNS neurons were not in a state of inflammation at 3 dpi with SARS-CoV-2. Surprisingly, inhibition of the JAK/STAT pathway resulted in high IL-6 protein levels in both mock- and SARS-CoV-2-infected CNS neurons (Figure 2I). Altogether, SARS-CoV2 infected CNS neurons with low efficiency and did not elicit a robust innate immune response.

### SARS-CoV-2 does not induce endoplasmic reticulum (ER) stress nor neurodegeneration in iPSC-derived CNS neurons

The UPR pathway controls ER homeostasis and becomes activated upon induction of ER stress. Chronic activation of the UPR can lead to neurodegenerative diseases (Pitale et al., 2017; Read and Schroder, 2021; Zhu and Lee, 2015). We investigated whether SARS-CoV-2 infection induced ER stress in CNS neurons by determining the expression level of activating transcription factor 4 (*ATF4*), CCAAT/enhancer-binding protein homologous protein (*CHOP*), binding immunoglobulin protein (*BiP*) and 94-kilodalton glucose regulated protein (*GRP94*). There were no signs of ER stress, such as upregulation of *ATF4* and *CHOP*, neither downregulation of *BiP* upon infection of CNS neurons compared to the mock-treated cells (Figure 3A). Interestingly, BiP expression, both at the transcriptional and translational level, was significantly increased in the CNS neurons exposed to SARS-CoV-2 (Figure 3A,B). Moreover, both SARS-CoV-2 infection and Ruxolitinib treatment inhibited the expression of *GRP94* (Figure 3A). Overall, these results suggest that CNS neurons responded to SARS-CoV-2 without severe dysregulation of the UPR pathway.

**Figure 3:**
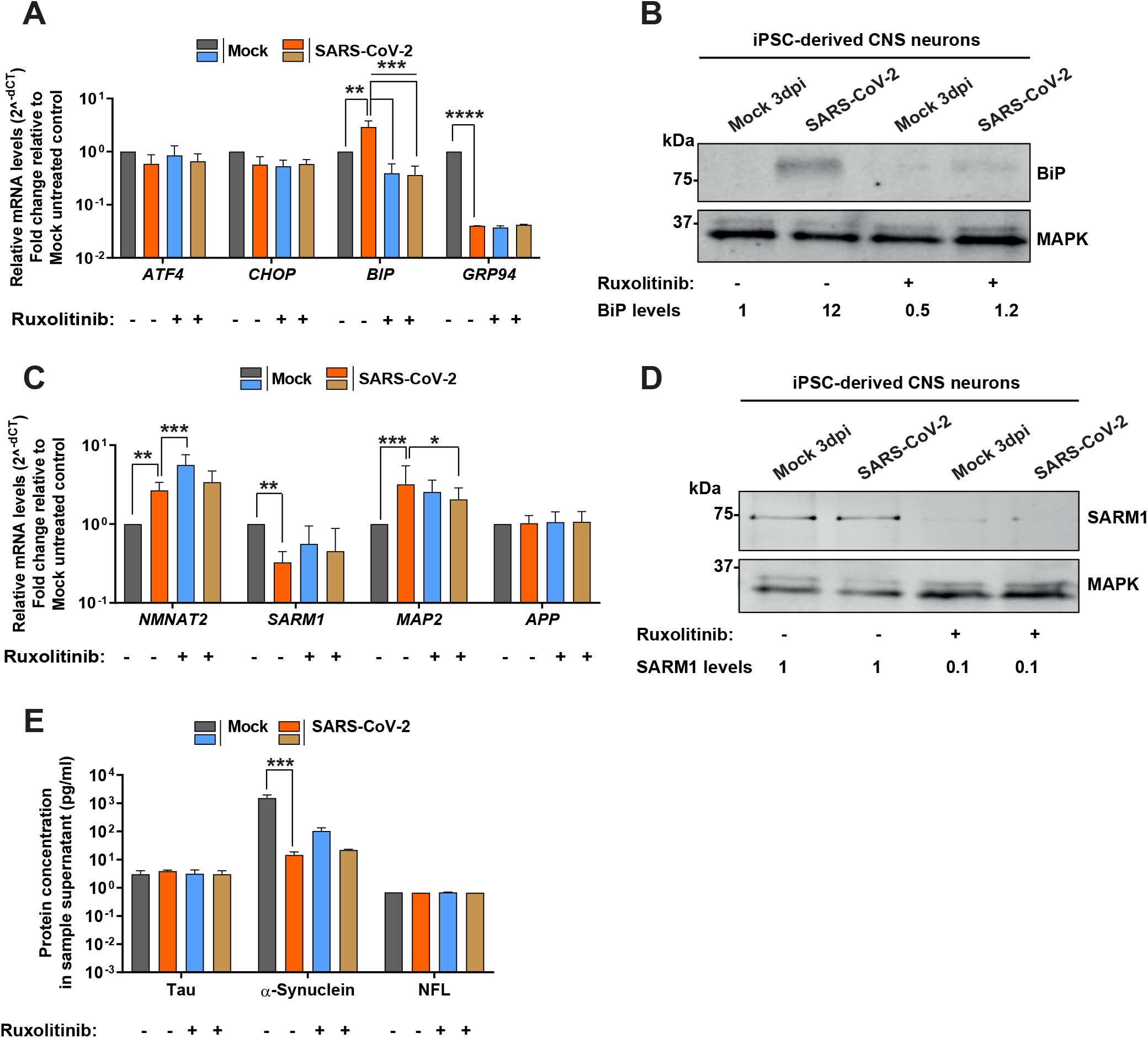
SARS-CoV-2 infected CNS neurons do not show cytopathic expression patterns. (A, C) Graphs showing relative expression of host genes in CNS neurons at 3 dpi. Gene expression was quantified by RT-qPCR and set relative to β-actin. Fold-change is relative to mock-infected, untreated control. (B, D) Immunoblots showing BiP (B), SARM1 (D) and MAPK (B,D) in cell lysates of CNS neurons obtained at 3 dpi. BiP and SARM1 levels were measured relative to MAPK using ImageJ. The numbers at the bottom of the blot represent the fold change to mock-treated control. Shown is one representative blot out of three independent ones. (E) Protein concentrations of released Tau, α-synuclein and Neurofilament L (NFL) determined by cytometric bead array multiplex analysis. In all graphs, error bars represent standard deviation of the arithmetic mean from three independent experiments. Statistical analyses for all experiments were determined using one-way ANOVA followed by the Dunnetts multiple comparison post-test. \**P*<0.03; \*\**P*<0.002; \*\*\**P*<0.0002; \*\*\*\**P*<0.0001; not significant comparisons are not indicated.

We also investigated the impact of SARS-CoV-2 infection on the expression of several genes involved in neurodegeneration such as *SARM1*, nicotinamide mononucleotide adenylyltransferase 2 (*NMNAT2*), β-amyloid precursor protein (*APP*) and the structural protein *MAP2*. CNS neurons infected with SARS-CoV-2 did not show patterns of axonal degeneration, but rather axonal survival, with increased transcript levels of *NMNAT2* and *MAP2*, reduced expression of *SARM1* and no changes in *APP* levels compared to the mock-treated CNS neurons (Figure 3C,D). Addition of Ruxolitinib further increased mRNA of *NMNAT2* in mock-infected cells and restored *SARM1* levels to those observed in mock-infected cells (Figure 3C). Interestingly, Ruxolitinib reduced SARM1 protein levels (Figure 3D). Ruxolitinib increased *MAP2* transcirpts in both mock- and SARS-CoV-2-infected cells compared to the samples without the inhibitor, while mRNA levels of *APP* did not change (Figure 3C). Moreover, the protein levels of tubulin associated unit (Tau) and neurofilament light chain (NFL) did not change upon infection and treatment with Ruxolitinib, while α-synuclein secretion was lower in the presence of the virus regardless of the presence of Ruxolitinib (Figure 3E).

Overall, we demonstrated that infection of human iPSC-derived CNS neurons with SARS-CoV-2 led to higher mRNA and protein expression of BiP, a central regulator of ER function (Hendershot, 2004). Moreover, we observed gene and protein expression profiles associated with neuronal survival and not with degeneration.

### SARS-CoV-2 infects iPSC-derived human PNS neurons and induces a type III IFN response

Infection of PNS neurons with SARS-CoV-2 has not been thoroughly studied. Therefore, we infected iPSC-derived PNS neurons with SARS-CoV-2 employing the same conditions as for the iPSC-derived CNS neurons. We observed similar NC protein amounts in the presence or absence of Ruxolitinib (Figure 4A). We also detected mRNA expression of *Membrane* and *NC*, which were reduced in the presence of Ruxolitinib (Figure 4B). The number of infected PNS neurons was higher than when we infected CNS neurons (Figure 4D). We also observed cytopathic effects in the cells, including lack of β-III-tubulin staining within some neurites, suggesting that neuronal degeneration took place (Figure 4D, yellow box). Neurons treated with Ruxolitinib had less cytopathic effect (Figure 4D). Moreover, there was less total β-III-tubulin protein upon SARS-CoV-2 infection compared to the mock-treated neurons, and a reversion to mock levels upon inhibition of the JAK/STAT pathway (Figure 4C). These results suggest that iPSC-derived PNS neurons undergo some degree of neurodegeneration upon SARS-CoV-2 infection, which is inhibited or delayed to some extent through suppression of the JAK/STAT pathway.

**Figure 4:**
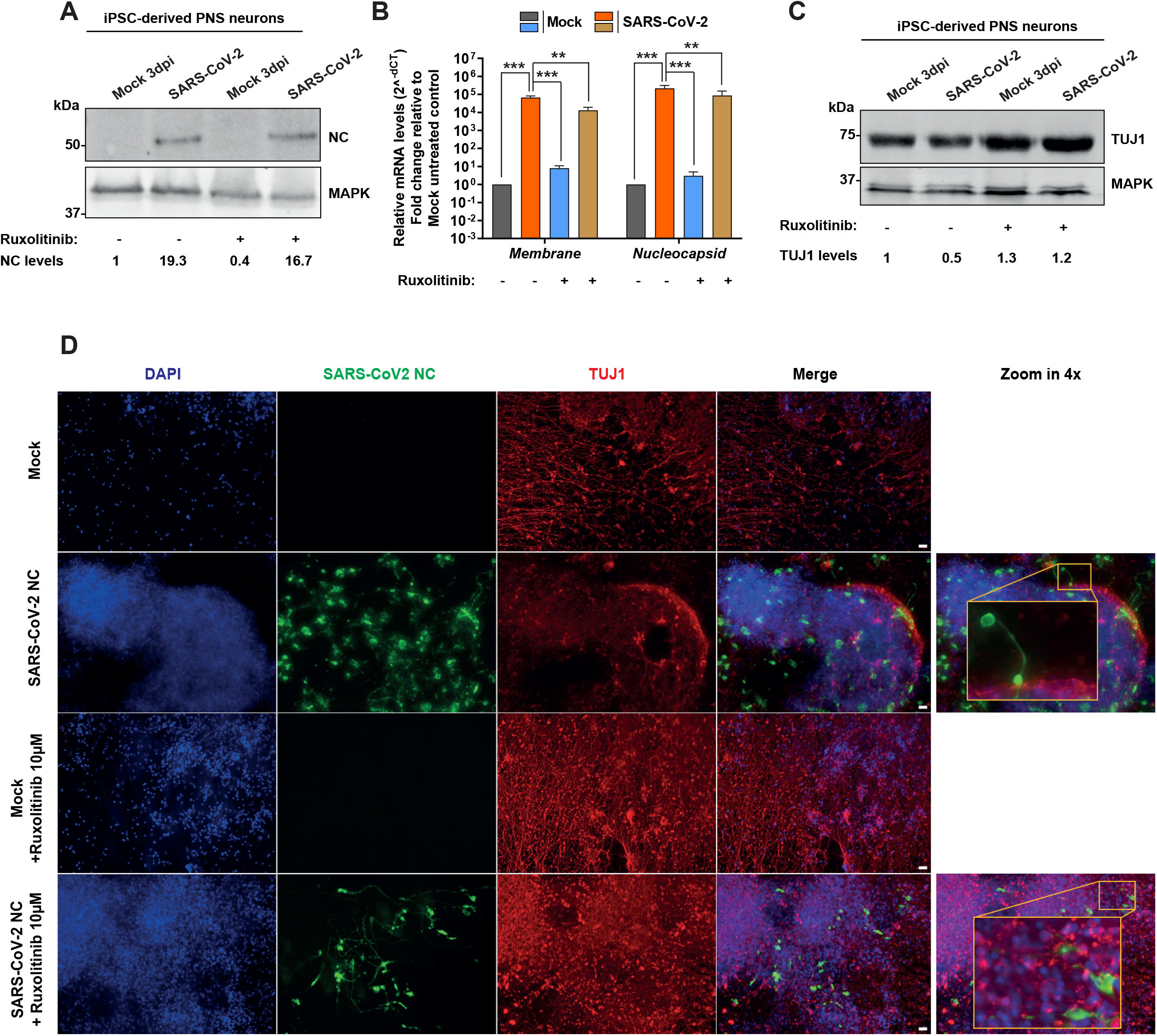
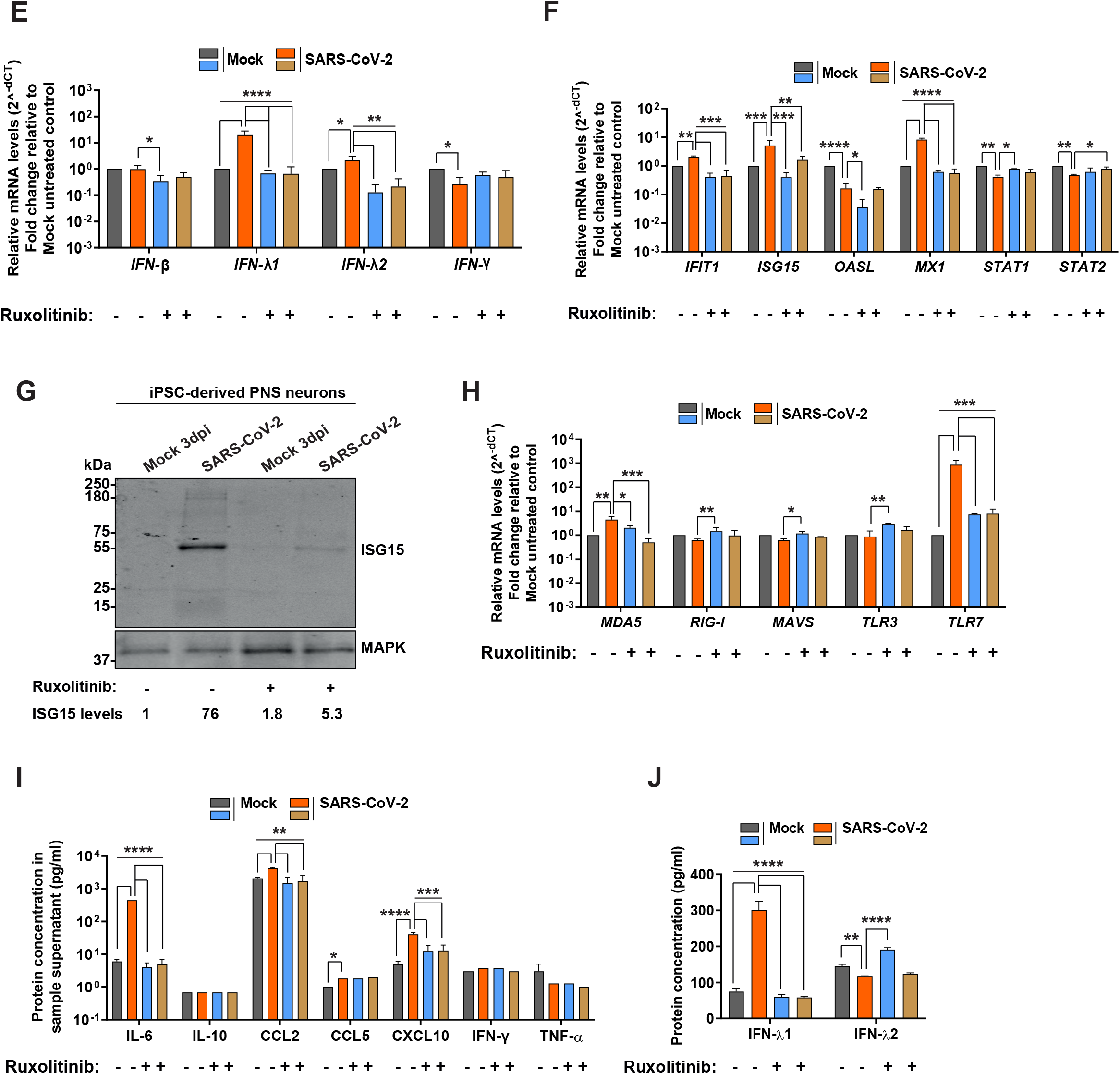
SARS-CoV-2 efficiently infects PNS neurons and induces robust type III IFN responses and ISG expression. (A) Immunoblot showing SARS-CoV-2 NC and MAPK in PNS neurons lysed at 3 dpi. NC levels were measured relative to MAPK using ImageJ. Shown is a representative blot from 3 independent experiments. (B) Graph showing relative expression of SARS-CoV-2 *Membrane* and *Nucleocapsid* mRNA in PNS neurons at 3 dpi. (C) Immunoblot showing TUJ1 and MAPK in cell lysates of PNS neurons at 3 dpi. TUJ1 levels were measured relative to MAPK using ImageJ. Shown is one representative blot out of three independent ones. (D) Immunofluorescence of PNS neurons fixed at 3 dpi and stained with anti-SARS-CoV-2 NC, anti-TUJ1 and DAPI. The right panels show a *zoom in* of 4x, applied using ImageJ. Scale bar = 100 μm; Amplification: 20X. (E, F, H) Graphs showing relative expression of host genes in PNS neurons at 3 dpi. (G) Immunoblot showing ISG15 and MAPK in lysates of PNS neurons at 3 dpi. ISG15 levels were measured relative to MAPK using ImageJ. Shown is a representative blot out of three independent ones. (I,J) Protein concentrations of released cytokines determined by cytometric bead array multiplex analysis (I) and ELISA (J) in the supernatant of infected PNS neurons. In all immunoblots, the numbers at the bottom of the blot represent the fold-change to mock-infected, untreated control. In all graphs showing gene expression, this was quantified by RT-qPCR and set relative to β-actin. Fold-change is relative to mock-infected, untreated control. In all graphs, error bars represent standard deviation of the arithmetic mean from three independent experiments. Statistical analyses for all experiments were determined using one-way ANOVA followed by the Dunnetts multiple comparison post-test. \**P*<0.03; \*\**P*<0.002; \*\*\**P*<0.0002; \*\*\*\**P*<0.0001; not significant comparisons are not indicated.

Next, we explored the effect of infection and inhibition of the JAK/STAT pathway in the expression of genes involved in the innate immune response during SARS-CoV-2 infection of PNS neurons. Interestingly, SARS-CoV-2 elicited a type III IFN response in PNS neurons, with significantly increased *IFN-λ1* and *IFN-λ2* gene expression and released IFN-λ1 protein, compared to mock-infected cells (Figure 4E,J). Addition of Ruxolitinib decreased the mRNA and protein levels of IFN-λ1 in SARS-CoV-2 infected cells, while it increased the protein amount of IFN-λ2 in the mock-infected cells (Figure 4E,J). Infection did not induce type I IFN and decreased slightly the expression of type II IFN, without affecting the protein level of IFN-γ (Figure 4E,I). We observed upregulation of *IFIT1, ISG15* and *Mx1* and downregulation of *OASL, STAT1* and *STAT2* mRNA levels in infected PNS neurons compared to mock-infected cells (Figure 4F). The increased ISG15 expression was also confirmed at the protein level (Figure 4G). The molecular weight of ISG15 was around 50 kDa, instead of 15 kDa, suggesting that it was ISGylated or associated with conjugates rather than the free form of ISG15 (Figure 4G). Ruxolitinib reduced ISG gene expression in the SARS-CoV-2-infected cells to basal levels (Figure 4F,G).

Next, we evaluated the expression levels of RNA sensors in PNS neurons upon SARS-CoV-2 infection. We did not observe significant changes in the mRNA levels of the analysed sensors with the exception of *MDA5* and *TLR7*, which were significantly higher in SARS-CoV-2-infected neurons compared to uninfected and to Ruxolitinib-treated cells, suggesting a relevance for these sensors in the recognition of SARS-CoV-2 in PNS neurons (Figure 4H). Analysis of the cytokine/chemokine profile showed an increase in IL-6, CCL2, CCL5 and CXCL10 in SARS-CoV-2 infected PNS neurons compared to mock-treated cells, revealing a phenotype that resembles the cytokine storm observed in severe COVID-19 patients (Figure 4I). Addition of Ruxolitinib inhibited the increase in cytokine expression observed during SARS-CoV-2 infection of PNS neurons (Figure 4I).

Overall, these results showed that SARS-CoV-2 infects more efficiently iPSC-derived PNS than CNS neurons, leading to higher innate immune response and loss of β-III-tubulin, which was less pronounced when the JAK/STAT pathway was inhibited.

### SARS-CoV-2 infection of PNS neurons results in expression of genes involved in the ER stress pathway and axonal degeneration

To study whether the activation of innate immune responses and the induction of stress pathways could lead to neurodegeneration, we infected PNS neurons with SARS-CoV-2 in the presence or absence of Ruxolitinib and analyzed the expression of genes related to the UPR pathway. Infection resulted in higher expression of *CHOP*, reduced mRNA levels of *GRP94* and no significant changes in transcription of *ATF4* and *BiP*, suggesting that PNS neurons were under stress conditions (Figure 5A,B). Treatment with Ruxolitinib decreased the expression of *CHOP*, and increased that of *GRP94*, supporting the hypothesis that the JAK/STAT pathway might play a role in the UPR in PNS neurons (Figure 5A).

**Figure 5:**
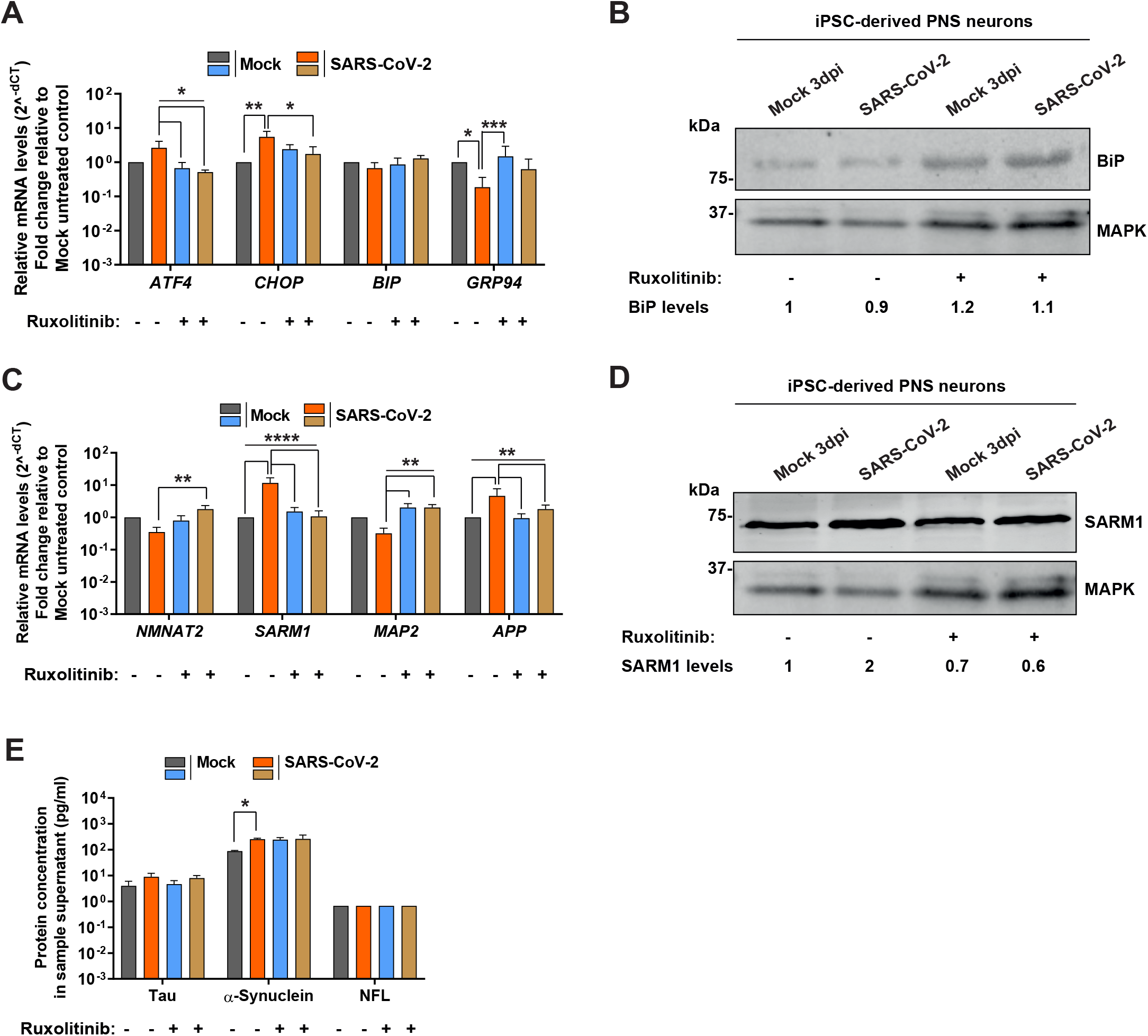
Stress response and axonal degeneration upon SARS-CoV2 infection of PNS neurons. (A, C) Graphs showing relative expression of host genes in PNS neurons at 3 dpi. Gene expression was quantified by RT-qPCR and set to β-actin. Fold-change is relative to mock-infected, untreated control. (B, D) Immunoblots showing BiP (B), SARM1 (D) and MAPK (B,D) in cell lysates of PNS neurons at 3 dpi. Relative BiP and SARM1 levels were measured using ImageJ. The numbers at the bottom of the blot represent the fold-change to mock-infected, untreated control. Shown is one representative blot from one experiment out of three independent ones. (E) Protein concentrations of released Tau, α-synuclein and Neurofilament L (NFL) determined by cytometric bead array multiplex analysis. In all graphs, error bars represent standard deviation of the arithmetic mean from three independent experiments. Statistical analyses for all experiments were determined using one-way ANOVA followed by the Dunnetts multiple comparison post-test. \**P*<0.03; \*\**P*<0.002; \*\*\**P*<0.0002; \*\*\*\**P*<0.0001; not significant comparisons are not indicated.

To further explore the reduced level of β-III-tubulin in PNS neurons upon SARS-CoV-2, we analysed the expression level of genes involved in axonal degeneration. *SARM1* mRNA and protein levels were significantly increased, pointing to its potential role in axonal degeneration in PNS neurons in the presence of SARS-CoV-2 (Figure 5C,D). SARM1 is essential and sufficient to trigger neurite degeneration (Figley et al., 2021; Gerdts et al., 2015; Osterloh et al., 2012). Additionally, *MAP2* transcript levels were reduced -although not significantly-, while those of *APP* were significantly increased in SARS-CoV-2-infected cells compared to the mock-infected cells (Figure 5C).

Interestingly, treatment with Ruxolitinib ameliorated the neurodegenerative phenotype by reverting the expression profile of *NMNAT2, SARM1, MAP2* and *APP*, suggesting a link between the innate immune response and neurodegeneration in PNS neurons infected with SARS-CoV-2 (Figure 5C,D). In addition, we observed significantly higher levels of released α-synuclein in the supernatant of PNS neurons infected with SARS-CoV-2 than in mock-infected neurons (Figure 5E).

In conclusion, our results indicate that CNS neurons were more resistant than PNS neurons to SARS-CoV-2 infection. They also suggest that a combination of direct effects of SARS-CoV-2 and innate immune mechanisms induce neuronal damage in iPSC-derived human PNS neurons.

## Discussion

Many COVID-19 patients suffer neurological complications. The protective role of the innate immune response of PNS and CNS neurons and its potential involvement in pathological outcomes upon SARS-CoV-2 infection are not well understood. Here, we addressed the role of the innate immune response upon SARS-CoV-2 infection of human iPSC-derived CNS and PNS. Our results indicate that infection of PNS neurons triggers IFN-λ1 expression that contributes to neuronal damage.

Infection of CNS neurons -with properties of striatal MSNs-was very inefficient and did not change after inhibition of the JAK/STAT pathway. These results are similar to those obtained in post-mortem human brains, which showed infection of cortical neurons, and lack of viral antigen in MSNs (Song et al., 2021). Exposure of CNS neurons to SARS-CoV-2 did not lead to a robust innate immune response, ER stress response and neurodegeneration when investigating changes within the total cell population, probably due to the inefficient infection. The expression of type I and III IFN was reduced, while that of type II IFN increased at the mRNA level. Another study reported induction of type III IFN and IL-8 during abortive infection of iPSC-derived CNS neurons (Bauer et al., 2021). Interestingly, despite the low number of infected CNS neurons, BiP was upregulated at the mRNA and protein levels. BiP is cytoprotective and may contribute to the lack of cytopathic effects observed in CNS neurons upon SARS-CoV-2 infection (Lewy et al., 2017). Moreover, there was also increased expression of NMNAT2, reduced levels of SARM1, higher level of the cytoskeletal protein β-III-tubulin and no increase in genes associated with neurodegeneration.

Interestingly, SARS-CoV-2 infected human iPSC-derived PNS neurons more efficiently than the CNS neurons, despite similar levels of ACE2, TMPRSS2 and Nrp-1. It is possible that SARS-CoV-2 infects PNS neurons through an ACE2-independent entry mechanism, as has been previously described for several cell lines and T lymphocytes (Hoffmann et al., 2022; Shen et al., 2022). Infection of PNS neurons correlated with the initiation of a type III IFN response, expression of *IFIT1, Mx1, ISG15* and pro-inflammatory cytokines like IL-6, CCL2, CCL5 and CXCL10 at 3 dpi. We did not detect induction of type I IFN responses under such conditions, probably because this response occurs earlier and is less sustained than the induction of type III IFN (Lazear et al., 2019).

The increased IFN-λ1 response in PNS neurons correlated with modification in expression of genes associated with the ER stress response (*CHOP* and *GRP94*), no changes in the master regulator of UPR (*BiP*), enhanced expression of neurodegeneration-related genes (*SARM1* and *APP*), an elevated level of proteins involved in neurodegeneration (SARM1 and α-synuclein) and lower levels of β-III-tubulin, essential protein maintaining the neuronal cytoskeleton. Studies performed in epithelial cells suggest a link between IFN-λ induction of SARM1 and UPR (Chen et al., 2021; Kanda et al., 2013). The observed increased expression in *CHOP* together with lack of changes in *BiP* expression could contribute to apoptosis and autophagy (Matsumoto et al., 2013) and thereby neurodegeneration in the human iPSC-derived PNS neurons. Interestingly, inhibition of the JAK/STAT pathway reverted this phenotype in PNS neurons, suggesting a link between the innate immune response, ER stress response, expression of SARM1 and neurodegenerative processes in the context of SARS-CoV-2 infection. Ruxolitinib also decreased SARS-CoV-2 gene expression, and this could also be linked to the lower cytopathic effect observed.

Neurons express IFN-λ and its receptor (Zhou et al., 2009). However, there are few reports on the role of the type III IFN response and its potential link to neurodegeneration during infection. The higher expression of IFN-λ1 in PNS neurons correlated with upregulation of *IFIT1, ISG15* and *Mx1*. ISGylation of ISG15 combined with high level of Mx1 were detected in peripheral mononuclear cells in symptomatic

COVID-19 patients (Schwartzenburg et al., 2022). Moreover, upregulation of ISG15 and its potential ISGylation could contribute to aberrant protein degradation promoting the accumulation of cellular proteins such as APP, and to increased autophagy, contributing to neurodegenerative diseases (Juncker et al., 2021).

SARM1 is both necessary and sufficient to initiate axonal degeneration (Gerdts et al., 2015; Osterloh et al., 2012), the first stage of many neurodegenerative diseases, including peripheral neuropathies (Geisler et al., 2016; Geisler et al., 2019). SARM1 is inactive in healthy axons but becomes active upon injury or insult, including infection, due to depletion of NMNAT2 (Figley et al., 2021). NMNAT2 promotes the synthesis of nicotinamide adenine dinucleotide (NAD+) through an enzymatic reaction of nicotinamide mononucleotide and adenosine triphosphate. SARM1 activation leads to hydrolysis of NAD into Nam, calcium influx and activation of calpains that degrade cytoskeletal proteins such as β-III-tubulin, causing neurite degeneration (Essuman et al., 2017; Horsefield et al., 2019; Posmantur et al., 1997; Yang et al., 2013). Upon infection of PNS neurons with SARS-CoV-2, there was a JAK/STAT-dependent activation of SARM1 that correlated with lower levels of β-III-tubulin and increase of neurodegenerative markers.

SARM1 plays different roles in the infection with different viruses. For instance, SARM1 knock out mice responded as wild type littermates to influenza, but were protected from neurodegeneration after CNS infection with vesicular stomatitis virus (Hou et al., 2013). This correlated with reduced levels of cytokine production in the CNS (Hou et al., 2013). On the contrary, mice lacking SARM1 expression were more susceptible to West Nile virus infection (Szretter et al., 2009). Our results also suggest a link between the innate immune response and SARM1 upon SARS-CoV-2 infection. The increased expression of SARM1 and reduced levels of cytoskeletal proteins upon SARS-CoV-2 infection of PNS neurons could lead to axonal degeneration and protection of the neuronal cell body from infection, as shown for Rabies virus (Sundaramoorthy et al., 2020). However, the degeneration of axons and nerve fiber loss could have pathological consequences, as observed in some COVID-19 patients (Abdelnour et al., 2020; Bitirgen et al., 2021; Oaklander et al., 2022).

Overall, we show that SARS-CoV-2 infected human PNS neurons more efficiently than CNS neurons and that the cytopathic effect leading to neurodegeneration was partly mediated by the initiation of an innate immune response. The pathological cytokine responses were characterized by high levels of IFN-λ1, expression of ISGs, cytokines, ER-stress genes and genes involved in triggering neurodegeneration, like SARM1, and accumulation of cellular proteins responsible for neuronal degeneration such as α-synuclein. Our model shows that blocking the JAK/STAT pathway ameliorates the neurodegeneration phenotype through decreasing *SARM1, ATF4, CHOP* and *APP* expression levels, reducing SARM1 protein level, as well as increasing β-III-tubulin and *MAP2*, linking the innate immune response and neurodegeneration.

To our knowledge, this is the first study indicating that type III IFN response contributes to neurodegeneration in human PNS neurons following infection. A considerable number of COVID-19 patients suffers from many symptoms associated with neuronal dysfunction in the PNS, including loss of innervation and increased neuropathy (Bitirgen et al., 2021; Oaklander et al., 2022). Our results cannot explain the broad range of different neurological symptoms from which COVID-19 patients suffer, but they provide a model to further explore the role of the infection and the innate immune response in the neuronal pathology and to test future preventive or therapeutic strategies.

## Materials and Methods

### Cells and viruses

Sensory neurons and MSNs were derived from iPSC and maintained as previously described (Kutschenko et al., 2021; Zhu et al., 2021). To differentiate sensory neurons, we employed small molecule derived neuronal precursor cells (smNPCs) generated from cord blood-derived iPSC (Naujock et al., 2014). The MSNs were differentiated from a control iPSC line (Schondorf et al., 2014). Vero76 cell line is a derivative of Vero cells, kidney epithelial cells from African green monkeys. Vero76 cells were cultured with DMEM supplemented with 10% foetal calf serum (FCS), 10 mM 4-(2-hydroxyethyl)-1-piperazineethanesulfonic acid (HEPES) and 1% of penicillin/streptomycin (Pen/Strep) in a humidified incubator at 37°C and with 5% CO_2_. SARS-CoV-2 β strain was kindly provided by Thomas F Schulz (Institute of Virology, Hannover Medical School, Germany). To prepare virus stocks, Vero76 cells were incubated with infection medium (DMEM supplemented with 2% FCS, 10 mM HEPES and 1% of Pen/Strep) containing SARS-CoV-2 β strain at an MOI of 0.01 plaque forming units (PFU)/cell for 48-72 hours in a humidified incubator at 37°C and with 5% CO_2_. Cell supernatant was collected and centrifuged at 450 g for 5 minutes. Virus aliquots were stored at -80°C. To determine the virus titre, we performed a plaque assay in VeroB4 cells, a derivative of Vero cells, kindly provided by Thomas F Schulz (Institute of Virology, Hannover Medical School, Germany), which were maintained in a humidified incubator at 37°C and with 5% CO_2._ Serial dilutions (10^−1^ to 10^−5^) were performed and a mixture of infection medium and virus was added to cells. 72 hours post infection, cells were washed and fixed with 10% formalin for 30 minutes. After evaporation of the formalin, 0.05% (w/v) of crystal violet in 20% methanol was added to each well and incubated during 30 minutes. The mixture was removed, cells were washed with distilled water and plaques were counted. The titre, defined as PFU/ml, was calculated using the formula PFU/ml = Number of plaques/infection volume*10^dilution^.

### Infection of iPSC-derived human neurons and Vero cells

To test for productive infection: 70 days differentiated MSNs were infected with SARS-CoV-2 β strain at an MOI of 0.01 to 1. 1 hour after addition of the virus (humidified incubator at 37°C and with 5% CO_2_) the inoculum was removed and 1:1 DMEM/F12 medium and Neurobasal medium supplemented with XN2, XB27 and 1% of Pen/Strep/Glutamine, 10 ng/ml of BDNF, 10 ng/ml GDNF, 25 ng/ml NGF and 50 μM dbcAMP was added to the cells. Cells were incubated in a humidified incubator at 37°C and with 5% CO_2_ for 24 and 48 hours; Vero76 cells were infected with SARS-CoV-2 β strain at an MOI of 0.01 (or with UV-inactivated SARS-CoV-2 β strain, kindly provided by Thomas Pietschmann, Experimental Virology, Twincore, Hannover) for 72 hours and maintained with DMEM supplemented with 2% FCS, 10 mM HEPES and 1% of Pen/Strep.

For the remaining experiments, differentiated CNS and PNS neurons were infected with SARS-CoV-2 β strain at an MOI of 0.01 for 72 hours, without removal of the inoculum after 1 hour. MSNs were incubated in 1:1 DMEM/F12 medium and Neurobasal medium supplemented with XN2, XB27 and 1% of Pen/Strep/Glutamine, 10 ng/ml BDNF, 10 ng/ml GDNF, 25 ng/ml NGF and 50 μM dbcAMP; PNS neurons were incubated in 1:1 DMEM/F12 medium and Neurobasal medium supplemented with 0.5% N2, 1% B27, 1% of Pen/Strep/Glutamine, 10 ng/ml BDNF, 10 ng/ml GDNF and 25 ng/ml NGF. To determine the role of the JAK/STAT pathway we added 10 μM of Ruxolitinib 18 hours prior to infection. Mock-infected cells were treated as infected ones without addition of the virus.

### RNA extraction and complementary DNA (cDNA) synthesis

Cells were harvested and total RNA was isolated using either the Nucleospin RNA mini kit (MACHEREY-NAGEL) or the Quick-DNA/RNA™ Viral kit (Zymo Research) according to the manufacturer’s instructions. SARS-CoV-2-infected and Mock-infected samples were treated with RNA Shield (Zymo Research) to preserve their genetic integrity and inactivate the virus. In all the samples, DNase treatment was performed according to the manufacturer’s instructions (RQ1 RNase-Free DNase, Promega). RNA samples were stored at -80°C. RNA quality was confirmed using a NanoDrop NOVIX DS-11 Spectrophotometer (Biozym). cDNA synthesis was performed using the LunaScript RT SuperMix Kit (NEB) according to the manufacturer’s instructions. cDNA was stored at -20°C. To avoid cell number bias, the same amount of RNA was used for all the samples to generate the cDNA used in the following steps.

### Reverse transcriptase quantitative polymerase chain reaction (RT-qPCR)

After cDNA synthesis, quantification of relative mRNA levels of target genes was performed using the Luna^R^ Universal qPCR Master Mix (NEB) according to the manufacturer’s instructions. Primer pairs (Sigma Aldrich) are listed in the table below. The RT-qPCR run was performed in a qTower3 (Analytic Jena) and the relative mRNA levels were obtained using the 2^-ΔCT^ method. The normalization for the NC and membrane levels was performed by setting an arbitrary CT value of 30 for the mock sample to reflect a lower detection limit and set mock to 1. *β-actin* was used as the internal reference gene. Each sample was analysed in triplicate and the data was analysed using the qPCRsoft 3.4 Software.

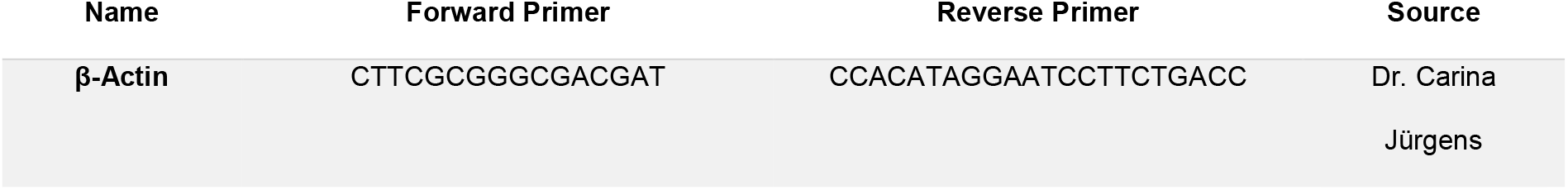
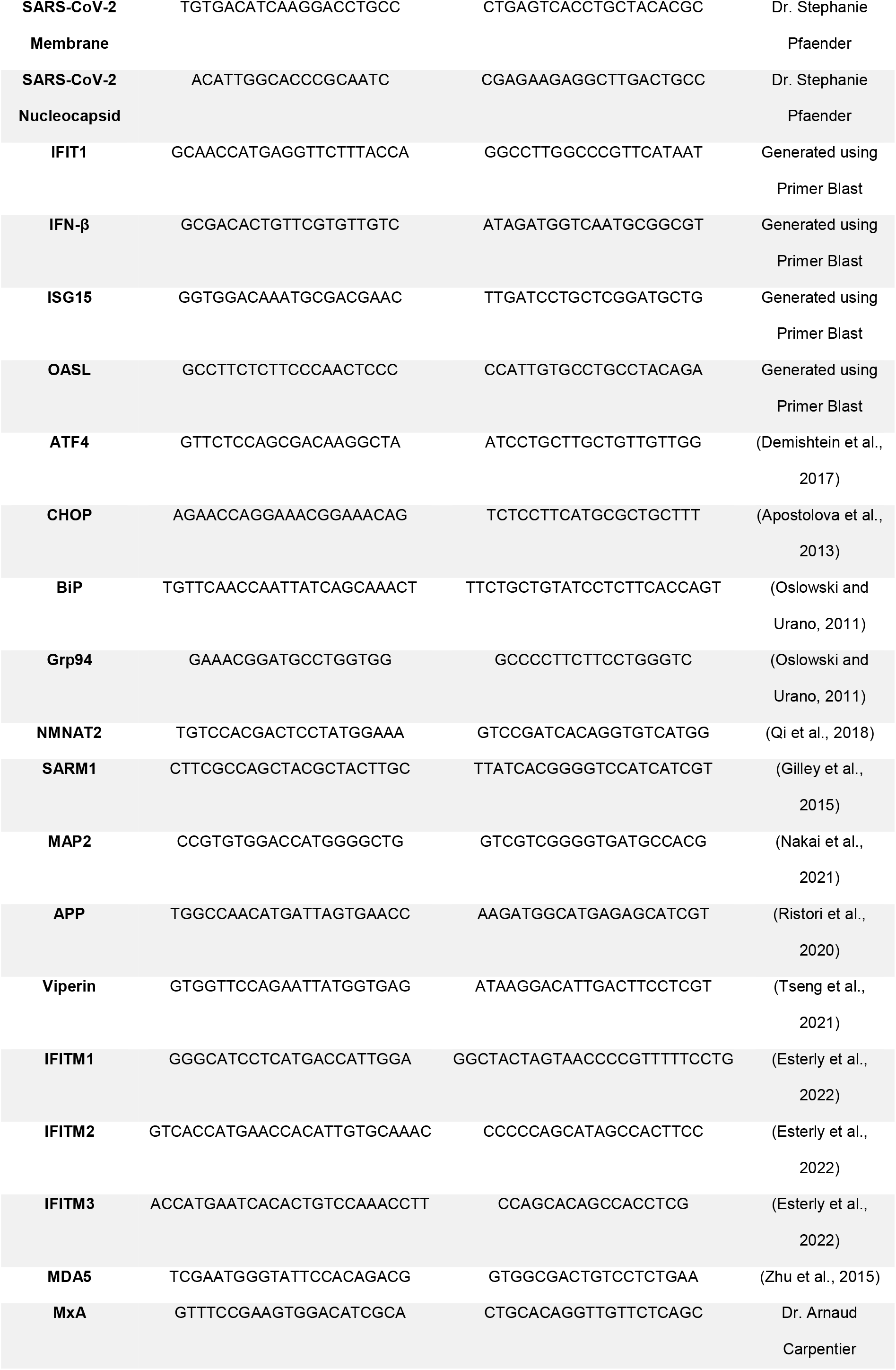
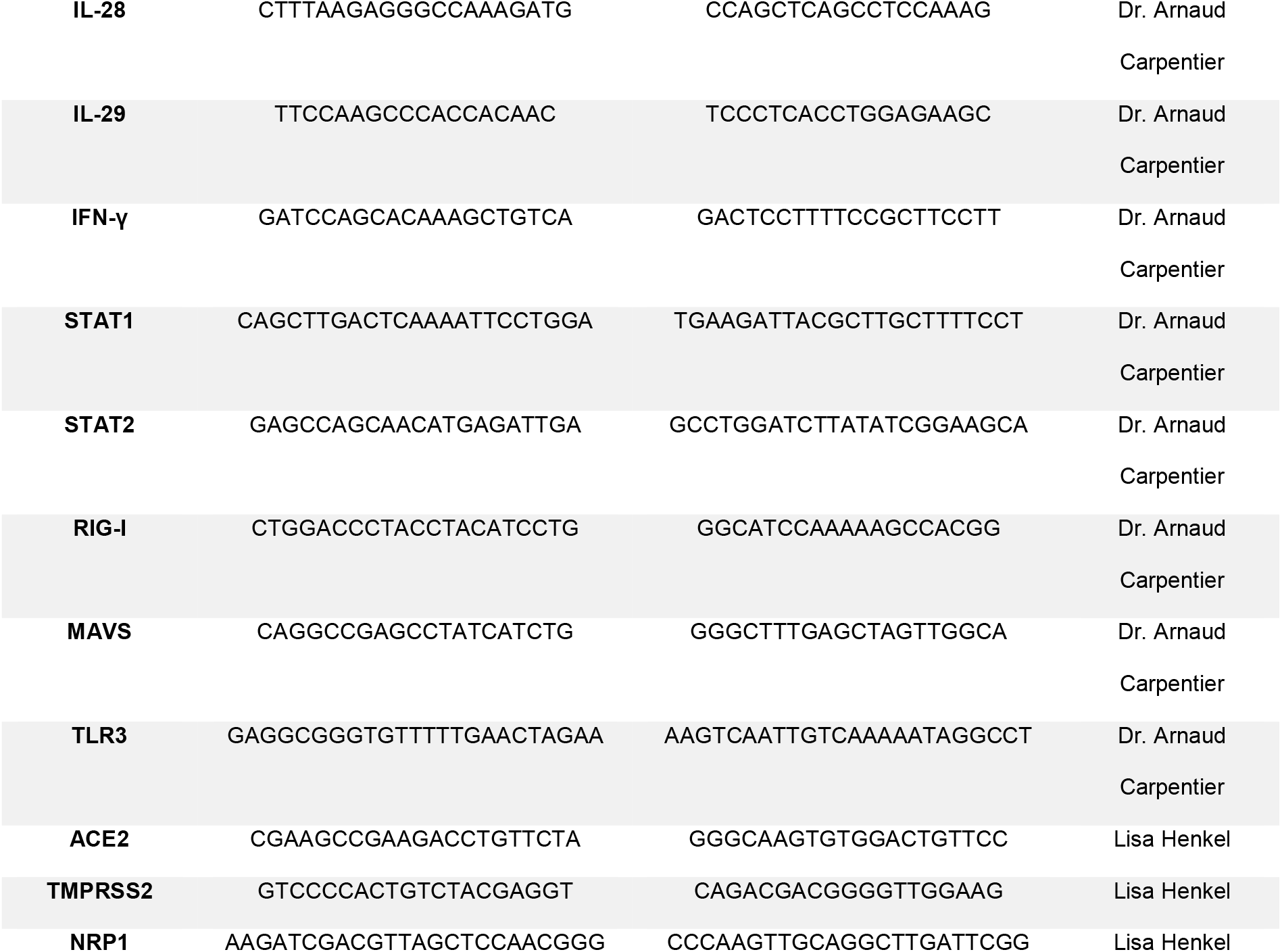

### Western blot

SARS-CoV-2- and mock-infected neurons, pre-treated with 10 μm Ruxolitinib or mock-treated, were lysed with Laemmli sample buffer and incubated at 95°C for 15 minutes to inactivate the virus. Proteins were run on a 10% sodium dodecyl sulfate-polyacrylamide gel electrophoresis and the transfer to a nitrocellulose membrane (Amersham™ Protran®) was performed in a wet system (Transfer chamber mini-Protean, BioRad). Membranes were blocked with 5% low fat powdered milk (Roth) and incubated overnight with the following primary antibodies: rabbit polyclonal anti-SARS-CoV-2 (COVID-19) nucleocapsid (1:5000; Biozol cat. No. GTX135357), rabbit polyclonal anti-ISG15 (1:1000; Cell Signalling Technology cat. No. 2743, a kind gift from Beate Sodeik), mouse monoclonal anti-β-III-Tubulin (Tuj1, 1:300; Millipore cat. No. MAB5564), rabbit polyclonal anti-BiP/GRP78 (1:1000; Abclonal cat. No. A0241), rabbit polyclonal anti-SARM1 (1:1000; Novus Biologicals cat. No. NBP1-77200), goat polyclonal anti-ACE (1:1000; R&D Systems cat. No. AF933-SP), mouse monoclonal anti-p44/42 MAPK ERK1/2 clone L34F12 (1:2000; Cell Signalling Technology cat. No. 4696S). Membranes were then subjected to secondary antibody incubations for 30 minutes: IRDye 800 CW goat anti-mouse IgG (Li-COR cat. No. 926-32210), IRDye 800 CW goat anti-rabbit IgG H+L (Li-COR cat. No. 925-32211), IRDye 680 RD donkey anti-mouse IgG (Li-COR cat. No. 925-68072), IRDye 680 RD goat anti-rabbit IgG H+L (Li-COR cat. No. 926-68071), IRDye 680 RD donkey anti-goat IgG (Li-COR cat. No. 925-68074). Detection was performed with the Li-COR Odyssey Imager system (Li-COR Biosciences) and quantification was done using ImageJ software. Protein levels were calculated using MAPK as the internal control and Mock untreated control was set to 1.

### Immunofluorescence

CNS and PNS neurons were differentiated on coverslips, treated with Ruxolitinib and infected with SARS-CoV-2 as detailed in methods section “Infection of iPSC-derived human neurons and Vero cells”. At 3 dpi, neurons on coverslips were fixed with 4% paraformaldehyde and washed with phosphate-buffered saline (PBS). Next, the cells were incubated with a blocking/permeabilization solution (2.5% bovine serum albumin – BSA, 5% goat serum, 0.3% Triton-X-100 in PBS) for 1 hour at room temperature. Neurons were stained with the following primary antibodies overnight at 4°C: mouse monoclonal anti-β-III-Tubulin (Tuj1, 1:1000; Millipore cat. No. MAB5564) and rabbit polyclonal anti-SARS-CoV-2 (COVID-19) nucleocapsid (1:1000; Biozol cat. No. GTX135357). Antibody dilutions were made in blocking/permeabilization solution. After washing, the cells were incubated with the following secondary antibodies: Alexa Fluor™ 633 Goat anti-Mouse IgG (H+L) Highly Cross-Adsorbed Secondary Antibody (1:500; Thermo Fisher Scientific cat. No. A21052), Alexa Fluor™ 647 Donkey anti-Mouse IgG (H+L) Highly Cross-Adsorbed Secondary Antibody (1:500; Thermo Fisher Scientific cat. No. A315712) and Alexa Fluor™ 488 Goat anti-Rabbit IgG (H+L) Highly Cross-Adsorbed Secondary Antibody (1:500; Thermo Fisher Scientific cat. No. A11034). Neurons were simultaneously stained with 4=,6-diamidino-2-phenylindole (DAPI) to detect the nuclei. After washing, the coverslips were mounted with ProLongTM Gold antifade reagent (Thermo Fisher Scientific, cat No. P36934). Images were obtained using a motorized Inverted Fluorescence Microscope Zeiss Axio Observer Z1 (Zeiss), and processed with ImageJ Sofware. Scale bar was set to 20 μm (CNS neurons) or 100 μm (PNS neurons). Images of CNS neurons were obtained with an amplification of 100X and PNS neurons of 20X with a zoom-in of 4x to better visualize the axons/dendrites.

### Multiplex assay

Supernatant (SN) of SARS-CoV-2- and mock-infected neurons, pre-treated with 10 μm Ruxolitinib or mock-treated, was collected. Virus was inactivated by incubating the SN with 7.5% of sodium bicarbonate on ice for 10 minutes. Then, SN was incubated with 0.1% of β-propiolactone (BPL) for 72 hours at 4°C. BPL was hydrolysed by incubating the SN at 37°C for 2 hours. SN was then stored at -20°C until use. To measure the cytokines/chemokines present in the SN we chose the LEGENDplex™ bead-based immunoassay and quantified the following cytokines/chemokines from the panels COVID-19 Cytokine Storm 1 (CCL2, CCL5, CXCL10, IFN-γ, IL-10, IL-6; Biolegend cat. No. 741088) and Human Neurodegeneration Biomarkers 1 (NFL, Tau, α-synuclein; Biolegend cat. No. 741197). The assay was performed following the manufacturer’s instructions. We used the SN without applying any dilution factor. Data acquisition was achieved as advised in the manufacturer’s manual and analysis was performed using the LEGENDplex™ Data Analysis Software. Protein concentration (pg/ml) was measured as the predicted concentration relative to the standards of each panel. Error bars represent standard deviation of the arithmetic mean from three independent experiments.

### Enzyme-linked immunosorbent assay (ELISA)

Supernatant (SN) of SARS-CoV-2 and mock-infected neurons, pre-treated with 10 μm Ruxolitinib or mock-treated, was collected and inactivated as mentioned in the Methods section “Multiplex assay”. To measure the protein concentration of IFN-λ1 and -λ2 present in the SN we performed an ELISA following the protocol from the manufacturer: IFN-λ1 (IL-29) from Invitrogen/Thermo Fisher Scientific cat. No. 88-7296-22; IFN-λ2 (IL-) from Invitrogen/Thermo Fisher Scientific cat. No. 88-52102-22. Washing buffer (PBST 0.05% Tween 20 (Sigma-Aldrich cat. No. P1379-25ML)) and stop solution (0.16 M Sulfuric Acid (Merck cat. No. 7664-93-9)) were prepared in-house. Absorbance values for IFN-λ1 and -λ2 were measured at 450 nm and protein concentration (pg/ml) was calculated using a standard curve created using Microsoft Excel.

### Statistical analysis

The significant value (*P* value) was calculated using GraphPad Prism by performing one-way ANOVA followed by Dunnetts’ multiple comparison post-test. Each gene was analysed separately and group comparisons were performed. In most of the cases Mock untreated control was set as 1. For the most of the data, logarithmic transformation was performed to reinsure Gaussian distribution. Error bars represent standard deviation of the arithmetic mean from three independent experiments. Statistical significance was shown as * *P*<0.03; \*\**P*<0.002; \*\*\**P*<0.0002; \*\*\*\**P*<0.0001; ns, not significant.

## Acknowledgements

We thank Stephanie Pfänder (Ruhr University Bochum) for scientific advice and protocols. We are thankful to Saskia Stein and Amelie Wachs (Institute of Virology, Hannover Medical School, Germany) for help during the introduction at the BSL3 laboratory. We thank Arnaud Carpentier (Twincore, Hannover), Carina Jürgens (Institute of Virology, Hannover Medical School, Germany) and Stephanie Pfänder for providing primer sequences. We are thankful to Thomas F. Schulz (Institute of Virology, Hannover Medical School, Germany) for providing the SARS-CoV-2 β strain and to Thomas Pietschmann for providing the UV-inactivated virus. We thank Thomas F. Schulz and Thomas Pietschmann for the Vero cells. We thank Beate Sodeik (Institute of Virology, Hannover Medical School, Germany) for providing the anti-ISG-15 antibody. We thank Ruth Knorr of the BSL3 facility at Hannover Medical School as well as Katja Branitzki-Heinemann and Maren von Köckritz-Blickwede of the BSL3 facility at University of Veterinary Medicine Hannover for their continuous support. The authors have no conflicting financial interests.

