## Supplemental Figures for "Innate immune response to SARS-CoV-2 infection contributes to neuronal damage in human iPSC-derived peripheral neurons"

### Supplemental information:

**Supplementary Figure 1**: SARS-CoV-2 infects CNS neurons and Vero76 cells. (A) Western blot showing detection of SARS-CoV-2 NC protein and MAPK in lysates of Vero76 cells infected with SARS-CoV-2 β strain, UV-inactivated SARS-CoV-2 or mock-infected for 48 hours. Shown is a representative blot from 3 independent experiments. (B,C) Graphs showing relative expression of *Membrane* and *Nucleocapsid* mRNA (left and right panels, respectively) detected by RT-qPCR in CNS neurons infected with SARS-CoV-2 at an MOI of 0.01, 0.1 and 1 or mock infected for 1 hour, 24 hours and 48 hours. Gene expression was set relative to β-actin and fold-change is relative to mock control.

**Supplementary Figure 2**: SARS-CoV-2 does not elicit a type II IFN response in CNS and PNS neurons. (A,B) Immunoblot to detect IFN-γ and anti-MAPK in lysates of CNS and PNS neurons at 3 dpi. Shown are representative blots from 3 independent experiments.

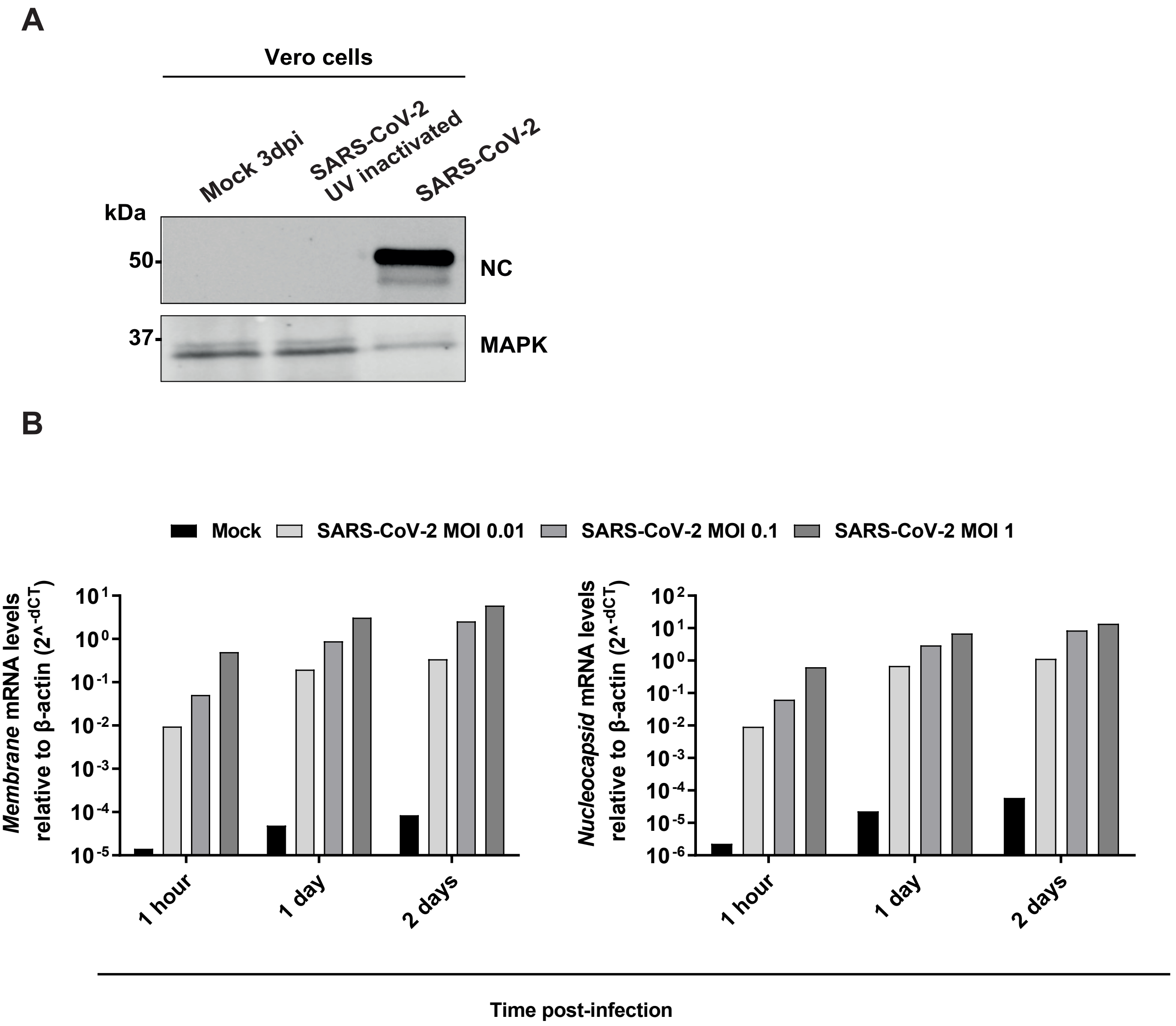
A
Vero cells
Mock 3dpi
SARS-CoV-2 UV inactivated
SARS-CoV-2
kDa
50
37
NC
MAPK
B
Mock
SARS-CoV-2 MOI 0.01
SARS-CoV-2 MOI 0.1
SARS-CoV-2 MOI 1
Membrane mRNA levels relative to β-actin (2^-dCT)
Nucleocapsid mRNA levels relative to β-actin (2^-dCT)
1 hour
1 day
2 days
Time post-infection

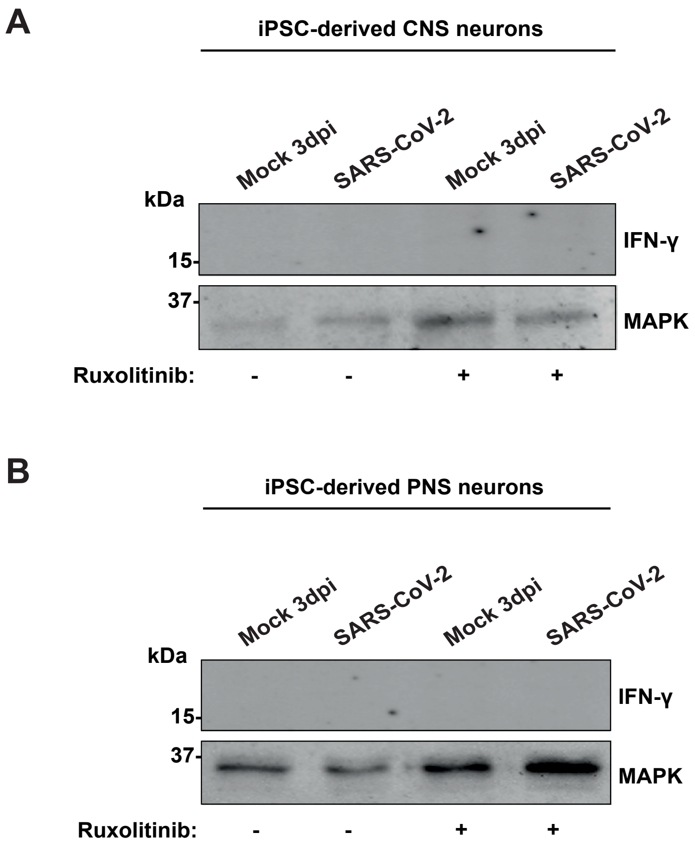
A
iPSC-derived CNS neurons
Mock 3dpi
SARS-CoV-2
Mock 3dpi
SARS-CoV-2
kDa
15-
37-
IFN-γ
MAPK
Ruxolitinib:
-
-
+
+
B
iPSC-derived PNS neurons
Mock 3dpi
SARS-CoV-2
Mock 3dpi
SARS-CoV-2
kDa
15-
37-
IFN-γ
MAPK
Ruxolitinib:
-
-
+
+
